# Cell-Intrinsic Vulnerability and Immune Activation Cooperate to Drive Degeneration in a Mitochondrial Complex I Deficiency Model of Optic Neuropathy

**DOI:** 10.1101/2025.10.08.681245

**Authors:** Daniela Santamaría-Muñoz, Raenier V. Reyes, Miranda R. Krueger, Andrea García-Llorca, Brennan Marsh-Armstrong, Xin Duan, Yang Hu, Derek S. Welsbie, Nicholas Marsh-Armstrong, Elisenda Sanz, Albert Quintana, Sergi Simó, Anna La Torre

**Affiliations:** Department of Cell Biology and Human Anatomy, University of California Davis. Davis, CA, USA; Department of Anesthesiology, Maine Health, Maine Medical Center, ME, USA; Department of Ophthalmology, University of California San Francisco. San Francisco, CA, USA; Department of Ophthalmology. Stanford University. Palo Alto, CA, USA; Department of Ophthalmology, University of California San Diego. San Diego, CA, USA; Department of Cellular Biology, Physiology and Immunology. Universitat Autònoma de Barcelona. Bellaterra, Barcelona, Spain

## Abstract

Mitochondrial dysfunction is a central hallmark of many optic neuropathies, yet the mechanisms linking intrinsic metabolic stress to retinal ganglion cell (RGC) degeneration remain unclear. To bridge this gap, we developed conditional transgenic models targeting the mitochondrial complex I subunit *Ndufs4* in the retina. Broad deletion of *Ndufs4* in the retina resulted in vision loss, progressive RGC degeneration, and pronounced immune activation before overt RGC death. Strikingly, depletion of myeloid cells significantly preserved RGCs, demonstrating that inflammation is not simply a downstream consequence but a participant in the degeneration process. To further distinguish between intrinsic and extrinsic mechanisms, we generated a mosaic model in which only subsets of retinal cells lacked *Ndufs4*. In this paradigm, the degeneration first appeared selectively in mutant regions, suggesting that mitochondrial impairment within RGCs is necessary to initiate vulnerability. At later stages, however, the degeneration extended beyond mutant territories, highly suggestive of a propagation through non-cell autonomous processes. Together, these findings support a model in which mitochondrial dysfunction creates the conditions for neuronal vulnerability, while immune responses govern the timing and extent of cell loss. This framework explains the consistent co-occurrence of metabolic deficits and neuroinflammation in optic neuropathies and highlights the importance of their interactions in disease progression. By clarifying the intersection of intrinsic and extrinsic mechanisms, this work advances our understanding of RGC degeneration and provides a conceptual basis for deciphering pathogenic processes across diverse optic neuropathies.

## Introduction

Optic neuropathies, including glaucoma, Leber’s hereditary optic neuropathy (LHON), and dominant optic atrophy (DOA), are leading causes of irreversible blindness worldwide [1–4]. A unifying trait of these conditions is the degeneration of retinal ganglion cells (RGCs), the projection neurons that convey visual information from the retina to the brain. RGCs are particularly susceptible to damage due to their unique anatomical structure, which includes unmyelinated segments within the retina, long axons, and high metabolic demands, making them especially vulnerable to disruptions in axonal transport and metabolic support [5–7]. Although mitochondrial dysfunction is increasingly recognized as a central pathogenic feature in most optic neuropathies [8–11], the mechanisms by which intrinsic metabolic stress drives progressive RGC degeneration and whether additional factors contribute to these processes remain incompletely understood.

Importantly, these metabolic challenges unfold within a retinal environment shaped by immune responses. Neuroinflammation consistently accompanies optic neuropathies in both animal models and human pathology [12–16], raising the possibility that immune mechanisms actively interact with metabolic stress to influence disease progression. In this direction, it has been proposed that both resident microglia and infiltrating monocytes actively participate in RGC pathology in different models [17, 18]. Microglia are among the earliest responders to stress and injury [13, 19]. Upon activation, these cells release inflammatory cytokines and reactive oxygen species and can engage in aberrant synaptic pruning and phagocytic activities, promoting further RGC dysfunction and death [20–23]. Yet microglia are not uniformly detrimental, several studies indicate that they can also contribute to tissue homeostasis and repair [24, 25]. For example, in optic nerve crush models, microglia have been implicated in debris clearance, which may support axon regeneration and reduce secondary damage [26], although some of these findings remain under debate [27].

In parallel, circulating monocytes can be recruited to the retina through disruption of the blood-retina barrier (BRB) or via permissive regions such as the optic nerve head (ONH), where the barrier is naturally attenuated [28]. Once in the tissue, these cells often adopt pro-inflammatory phenotypes and can exacerbate neurodegeneration [29–33]. Conversely, in some injury contexts, macrophage-derived factors can promote axon regeneration [34]. More recently, the adaptive immunity has also emerged as an important factor: CD4+ T cells infiltrate the retina and can contribute to RGC loss, while other studies have suggested that regulatory T cells could exert protective effects by limiting glial activation and promoting BRB integrity [35–39].

Taken together, these observations illustrate that the immune responses during optic nerve degeneration are complex and context- and timing-dependent. Based on current evidence, several non-mutually exclusive possibilities can be considered: immune responses may help mitigate ongoing degeneration by clearing debris and supporting tissue integrity; they may be required for pathological progression; or they could amplify the initial insult, triggering bystander degeneration of otherwise healthy cells. Distinguishing how these roles interact with intrinsic vulnerability is essential for understanding the progression of RGC degeneration.

The importance of context also underscores the need to investigate the roles of these immune responses within various disease models, as the initiating insult can influence the nature of inflammation. Thus, whereas ocular hypertension and optic nerve injury models primarily engage mechanical or traumatic pathways, mitochondrial dysfunction models uniquely enable direct investigation of how intrinsic metabolic stress shapes inflammation. These models therefore provide a powerful opportunity to disentangle how neuronal-intrinsic deficits interact with immune mechanisms to drive RGC degeneration, offering insights not accessible in other injury-based paradigms.

The mitochondrial complex I subunit Ndufs4 is a critical component of the electron transport chain, and its loss impairs mitochondrial function and neuronal viability [40, 41]. Several models of *Ndufs4* ablation have been reported [40–42], and these consistently exhibit retinal degeneration phenotypes [43–45]. However, many of these lines also have very short lifespans, which hinders the study of progressive neuronal degeneration. To circumvent this limitation, we have employed conditional knockout strategies to target *Ndufs4* in the retina, thereby modeling mitochondrial dysfunction in optic neuropathy without affecting other tissues. In addition to mice with widespread retinal deletion, we have also generated a mosaic model in which *Ndufs4* is ablated in defined retinal regions, while other areas retain wild-type expression, producing a stereotyped pattern of cellular *Ndufs4* loss. This design offers a unique platform to assess how intrinsic metabolic deficits render RGCs vulnerable and how extrinsic immune mechanisms contribute to the propagation of degeneration across retinal regions.

Using these models, we first examined the temporal and spatial dynamics of neurodegeneration and inflammation. Degeneration always began in regions lacking *Ndufs4*, indicating that intrinsic mitochondrial dysfunction is the primary driver. Prior to measurable RGC soma loss, myeloid cells became activated, establishing inflammation responses an early feature of the degenerative process. Interestingly, this activation occurs in both mutant and non-mutant regions, even though RGC loss remained confined to areas with Ndufs4 deficiency. Notably, depletion of myeloid cells was neuroprotective, indicating that the early innate immune activation is required for the degeneration in vulnerable regions. At later stages, degeneration spread into neighboring areas that retained Ndufs4, suggesting that additional mechanisms contributed to the late-stage disease progression. Together, our findings support a multi-step model in which mitochondrial dysfunction primes RGCs for degeneration, early myeloid-driven inflammation accelerates damage, and later-stage non-cell-autonomous processes further influence the course of the disease. These insights provide a framework for understanding how intrinsic and extrinsic factors converge to drive optic neuropathies and may inform strategies to prevent or slow vision loss.

## Materials and methods

### Animals

The Rax-Cre BAC transgenic line, Tg(Rax-Cre)NL44Gsat/Mmucd, was developed by the GENSAT project and cryopreserved at the Mutant Mouse Resource and Research Center (MMRRC), University of California, Davis (stock number: 034748-UCD; [46]). The α-Cre transgenic line was originally generated by [47], and the *Ndufs4* conditional allele was described by [41]. All animals were maintained on a mixed CD-1 × C57BL/6 genetic background. Genotyping was performed by PCR using protocols provided by The Jackson Laboratory (JAX) or MMRRC.

All animal procedures were approved by the University of California, Davis Institutional Animal Care and Use Committee (IACUC) and carried out in accordance with National Institutes of Health (NIH) guidelines. For experiments involving myeloid cell depletion, animals were fed either a standard AIN-76A Rodent Diet or AIN-76A supplemented with 1,200 ppm PLX5622 (MedChemExpress) beginning at P21 and continuing until tissue collection at P45. Both diets were prepared by Research Diets, Inc.

### Optomotor response

Visual function was assessed using optokinetic tracking with the OptoMotry system (Cerebral Mechanics) as previously described [48]. Mice were placed on an elevated platform surrounded by four monitors displaying rotating vertical sine wave gratings (100% contrast; 12 degrees/second). Spatial frequency was adjusted using a staircase method, increasing progressively until the experimenter could no longer detect a head-tracking response consistent with the direction of the rotating gratings.

### Immunohistochemistry

To obtain retinal cross-sections, enucleated eyes were fixed overnight at 4 °C in modified Carnoy’s fixative, consisting of 60% ethanol, 30% formaldehyde, and 10% acetic acid. Following fixation, tissues were embedded in paraffin blocks as described before [49]. Serial sections were cut at a thickness of 5 µm.

Paraffin-embedded tissue sections were deparaffinized and rehydrated. Permeabilization was performed for 15 minutes using 0.3% Triton X-100 in PBS. Next, antigen retrieval was carried out by heating sections in 0.01 M sodium citrate buffer (pH 8.0) at 95 °C for 5 minutes, repeated twice. This was followed by a 1-hour incubation in 0.5% Triton X-100 containing 2 N HCl. Sections were then incubated in a blocking solution consisting of 10% normal donkey serum in 0.1% Triton X-100 in PBS for 1 hour at room temperature. Primary antibodies (see Table 1 for details) were diluted in the blocking solution and applied to the sections for overnight incubation at 4 °C. After washing with PBS, sections were incubated for 1 hour at room temperature with species-specific, Alexa Fluor secondary antibodies, also diluted in blocking solution. Nuclear counterstaining was performed using 4′,6-diamidino-2-phenylindole (DAPI; Sigma-Aldrich) at a concentration of 1 μg/ml for 15 minutes. Finally, sections were washed and mounted using Fluoromount-G (Southern Biotech). Samples were imaged using an Olympus Fluoview FV4000 confocal microscope.

**Table 1.**
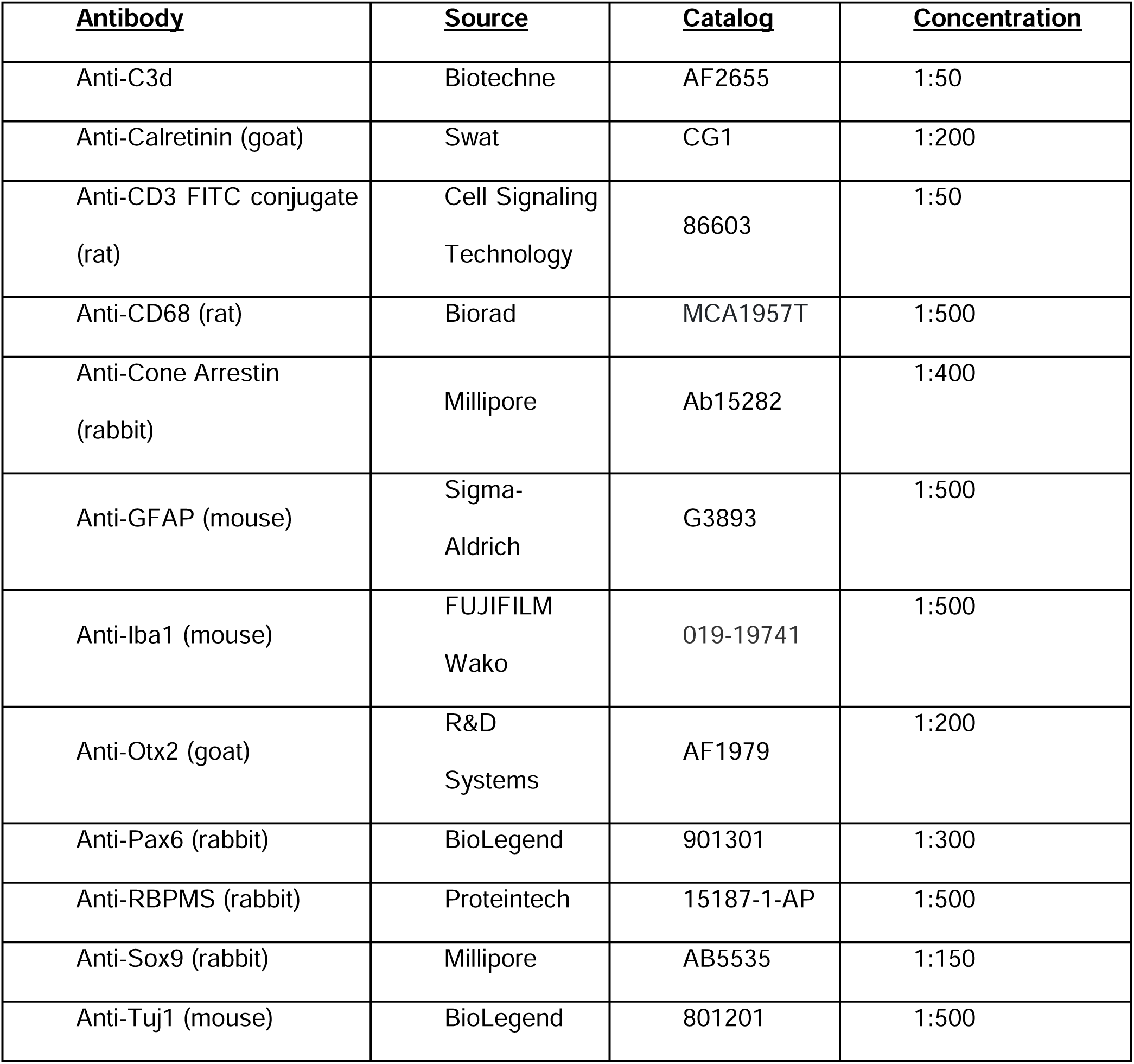
Primary antibodies.

| <u>Antibody</u> | <u>Source</u> | <u>Catalog</u> | <u>Concentration</u> |
| --- | --- | --- | --- |
| Anti-C3d | Biotechne | AF2655 | 1:50 |
| Anti-Calretinin (goat) | Swat | CG1 | 1:200 |
| Anti-CD3 FITC conjugate (rat) | Cell Signaling Technology | 86603 | 1:50 |
| Anti-CD68 (rat) | Biorad | MCA1957T | 1:500 |
| Anti-Cone Arrestin (rabbit) | Millipore | Ab15282 | 1:400 |
| Anti-GFAP (mouse) | Sigma-Aldrich | G3893 | 1:500 |
| Anti-Iba1 (mouse) | FUJIFILM Wako | 019-19741 | 1:500 |
| Anti-Otx2 (goat) | R&D Systems | AF1979 | 1:200 |
| Anti-Pax6 (rabbit) | BioLegend | 901301 | 1:300 |
| Anti-RBPMS (rabbit) | Proteintech | 15187-1-AP | 1:500 |
| Anti-Sox9 (rabbit) | Millipore | AB5535 | 1:150 |
| Anti-Tuj1 (mouse) | BioLegend | 801201 | 1:500 |

For retinal flat-mounts, enucleated eyes were immersed in 4% paraformaldehyde (PFA) in PBS for 30 minutes at room temperature. Retinas were then dissected and subjected to an additional 30-minute fixation in 4% PFA. After fixation, tissues were permeabilized for 1 hour in PBS containing 1% Triton X-100. After 2 hours in blocking solution (10% normal donkey serum in 1% triton X-100 in PBS), samples were incubated with primary antibodies diluted in the blocking solution and left overnight at 4°C. The next day, tissues were washed three times with PBS, then incubated with secondary antibodies also diluted in blocking solution for an additional overnight period at 4°C. Finally, retinas were washed in PBS and mounted using Fluoromount-G (SouthernBiotech) for imaging. Flat-mounts were imaged at 20x using an Axio Imager.M2 with Apotome.2 Microscope System (Zeiss), images were tiled to generate whole-retina composites.

### Analysis of RGC survival

RGCs were counted using an automated Python-based analysis pipeline described previously [50]. In brief, absolute background subtraction, morphological opening, and kernel-based background subtraction was used to eliminate sub-cellular-sized background noise. Then, repeated kernel-based blurring and erosion was used to separate individual cells from clusters. Adaptive thresholding and Euclidean distance mapping were used to binarize and then map object centers, with subsequent filtering for size and intra-center distance generating a list of coordinates for all objects likely representing a cell. This counting approach was demonstrated previously to vary from manual counting of retinal flat mount cells by less than 4 per 1000 cells. Total RBCs per retina were counted from images in which the entire retina was masked with an outline generated in Fiji. Regional quantification of RGCs was performed by manually masking four square 380 × 380 µm² regions of the retina and subjecting that region to the abovementioned analysis.

### Hematoxylin and eosin (H&E) staining

Tissue was prepared as described before [49]. Briefly, tissue sections were deparaffinized in xylene and subsequently rehydrated through a graded ethanol series. Hematoxylin staining was performed for 7 minutes, followed by a brief rinse in 0.5% acid alcohol (2.5 ml hydrochloric acid in 500 ml 70% ethanol) and then immersion in ammonia water (0.5 ml ammonia hydroxide in 500 ml H_2_O). Sections were then counterstained with eosin (500 ml 75% ethanol, 3.75 ml glacial acetic acid, 2.5 g eosin, 1 g phloxine B) for 6 minutes. After staining, slides were dehydrated through graded ethanol solutions, cleared in xylene, and mounted using Permount mounting medium (Fisher Chemical).

### Assessment of axonal degeneration

Optic nerves were dissected and fixed overnight with 5% glutaraldehyde in 0.1 M sodium phosphate buffer. Tissues were then rinsed twice in 0.1 M sodium phosphate buffer for 30 minutes and placed in secondary fixative in 1% osmium tetroxide/ 1.5% potassium ferrocyanide for 3 hours. After rinsing with 0.1 M sodium phosphate, samples were dehydrated in graded ethanol concentrations. Tissues were then placed in propylene oxide twice for 15 minutes and pre-infiltrated in half resin/ half propylene oxide overnight. The next day, tissues were infiltrated in 100% resin (450 ml dodecenyl succinic anhydride, 250 ml araldite 6005, 82.5 ml Epon 812, 12.5 ml dibutyl phthalate, and 450 μl benzyldimethylamine) for 5 hours. Samples were embedded with fresh resin and polymerized at 65°C overnight.

For PPD staining, the embedded tissues were sectioned with a Leica EM UC6 ultramicrotome at a thickness of 450 nm. Sections were then stained with 2% PPD in 50% ethanol for 18 minutes, followed by a washing step involving submersion in 100% ethanol 20 times, and then mounted with Cytoseal 60 (Electron Microscopy Sciences). Images were obtained at 40x magnification by Axio Imager.M2 with Apotome.2 Microscope System (Zeiss). Optic nerve axons were then quantified manually by selecting 50 x 50 µm^2^ segments. Values were then normalized to a control to show the final data as an axon percentage.

Optic nerve morphology was assessed using transmission electron microscopy. Optic nerve sections of 70 nm were cut transversely on a Leica Ultracut microtome (Leica Microsystems). Samples were collected on copper grids, counterstained with uranyl acetate and lead citrate, and then examined using a FEI Thermo Scientific Talos L120C transmission electron microscope equipped with a CETA 16MP camera at 80 kV. Electron microscopy images were blinded prior to the evaluation. Investigators were not aware of the experimental groups during the image acquisition. The images were all the same size and same magnification (SA 2000X).

### Analysis of Iba1_+_ Cell Population

Composite whole-mount images were generated from retinas stained with Iba1 and Tuj1. Tuj1 staining served as a reference to determine the retinal layer in which images were acquired, specifically distinguishing between the ganglion cell layer/retinal nerve fiber layer (GCL/RNFL) and the inner plexiform layer (IPL). To quantify Iba1_+_ cells in each layer, four square regions of interest (630 × 630 µm²) were selected per retina, positioned either near the retinal edge (periphery) or adjacent to the optic nerve head (central region). Iba1_+_ cells within each region were manually counted.

To assess the spatial distribution of Iba1_+_ cells, 10 cells were randomly selected from both the peripheral and central regions of each retina. For each selected cell, the distance to its five nearest neighboring Iba1_+_ cells was measured using Fiji. These distances were averaged to yield a single value per region and biological replicate.

### Cytokine array

Retinas were homogenized in PBS with protease inhibitors (Millipore Sigma, catalog no. P8340), followed by the addition of Triton X-100 to a final concentration of 1%. Homogenized tissue was frozen at -70°C, thawed, and centrifuged at 10,000 x g for 5 min. After removing cellular debris, protein concentration was determined using Qubit Protein Quantification (Thermo Fisher Scientifc). Cytokine proteomic profiling was then performed following the manufacturer’s instructions (R&D Systems, catalog no. ARY006). For each time-point and experiment, polls of 4 eyes were used, and all samples were run in technical duplicates. Two biological replicates were used per time-point. Cytokine array films were scanned and files were saved in an uncompressed format (TIFF). Quantification was performed using ImageJ (NIH). Images were first converted to 8-bit grayscale. The “Analyze > Set Measurements” tool was used to record integrated density and mean gray values. Circular regions of interest (ROIs) of identical size were drawn around each dot, and the same ROI was applied across all samples to maintain consistency. A background ROI of the same size was placed in a blank area of the membrane, and its value was subtracted from each measurement. The resulting background-subtracted integrated density values were used for analysis. The values were then log10-transformed and visualized in a clustered heatmap using hierarchical clustering of cytokines (rows). Clustering was performed using Euclidean distance and average linkage. Heatmaps were generated in Python 3.11 with the seaborn package (v0.12.2).

### RNA scope *in situ* hybridizations

For *in situ* hybridazation, protocols were performed according to the RNAscope Multiplex Fluorescent Reagent Kit v2 Assay manual using the provided reagents and as described before [49]. In summary, after sample fixation and preparation described above, paraffin slides were baked at 60°C and subsequently deparaffinized. The tissue was pretreated with 5% hydrogen peroxide at RT for 10 min, target antigen retrieval for 20 min at 99°C, and protease plus for 30 min at 40°C. Slides were then incubated with the appropriate hybridization probes for 2 hr at 40°C. Probes include: Cxcl10 (408921-C2) and Cxcl9 (489341). Images were obtained by Fluoview FV4000 confocal microscope (Olympus).

### Statistics

The number of biological replicates and the statistical tests used for each experiment are specified in the corresponding figures and figure legends. Data are presented as mean ± SEM. Statistical significance was defined as *p* < 0.05. All analyses and data visualizations were carried out using GraphPad Prism 9.

## Results

### Conditional *Ndufs4* deletion in the retina causes vision loss and RGC degeneration

To establish a model of mitochondrial dysfunction in the retina while avoiding the limited survival observed in systemic *Ndufs4* knockout mice [41, 51] and other models [42], we utilized a conditional strategy. We crossed *Ndufs4* conditional knockout mice (Ndufs4^fl/fl^ [41]) with a Rax-Cre transgenic driver [46]. As reported before, Rax is expressed in the optic vesicle and all its derivatives, including all the cells in the retina and the retinal pigment epithelium (RPE) (Fig.1A, [52, 53]. Although *Ndufs4* is ablated in retinal progenitors from embryonic day 8, we did not observe retinal developmental defects, and these animals are healthy and exhibit normal lifespans.

**Figure 1.**
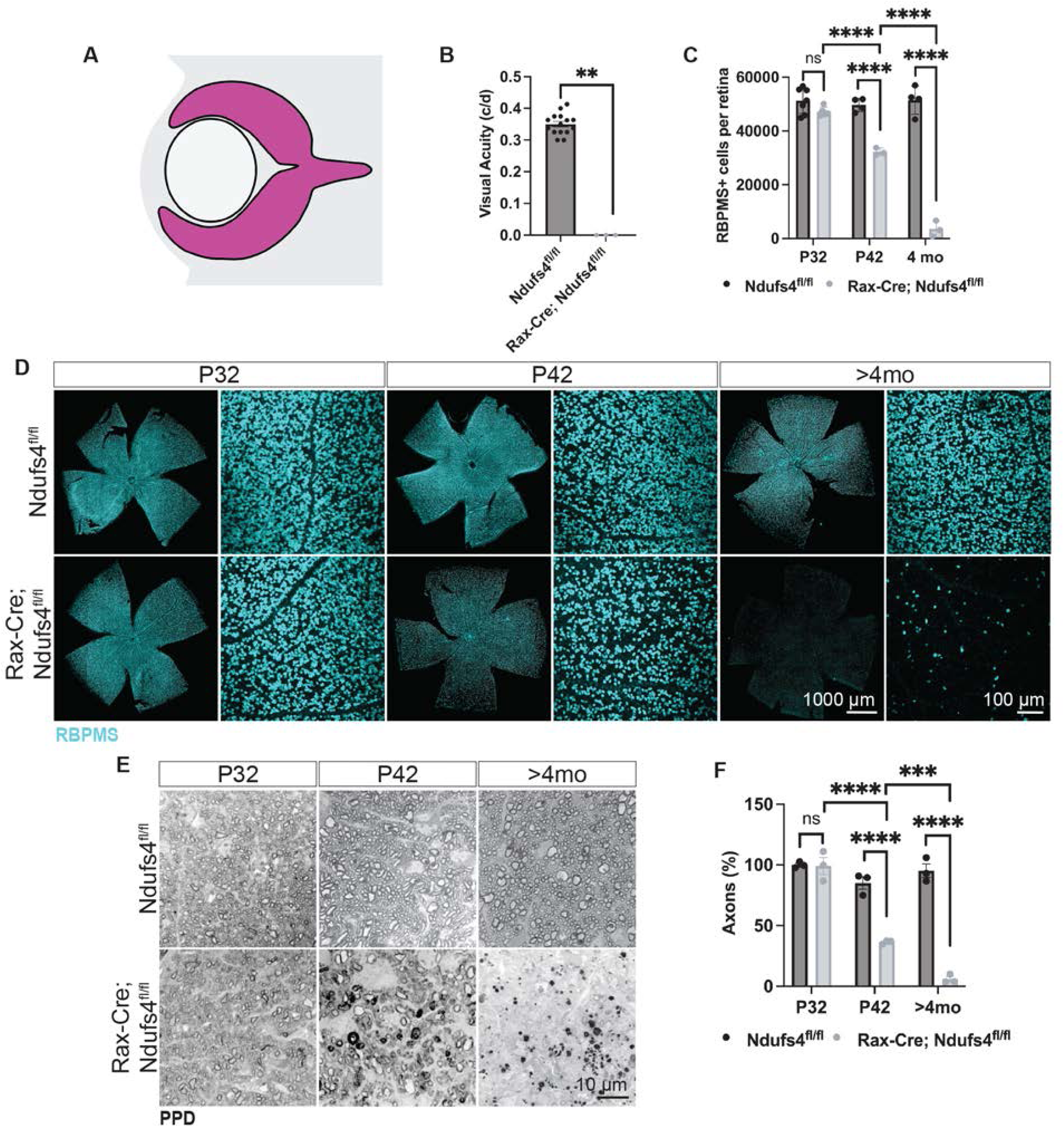
Rax-Cre; Ndufs4^fl/fl^ mice exhibited vision loss and progressive RGC degeneration. A) Simplified diagram of the embryonic retina showing the expression pattern of Cre (magenta) in Rax-Cre. B) Optomotor response of 4-month animals (Mann-Whitney U test, **P < 0.01). C) Quantification of RBPMS+ RGCs in flat-mounted retinas over time (two-way ANOVA, n.s. not significant and ****P < 0.0001). D Flat-mounted retinas labeled with RBPMS (teal) E) Paraphenylenediamine (PPD) staining to label myelinated axons in optic nerve sections demonstrates axonal loss in Rax-Cre; Ndufs4^fl/fl^ animals. F) Quantification of the number of axons in PPD-stained optic nerve sections (two-way ANOVA, n.s. not significant, ***P < 0.001, and ****P < 0.0001).

We then asked whether retinal function was preserved in adulthood. For all experiments, we compared Rax-Cre; Ndufs4^fl/fl^ mice with littermate controls (Ndufs4^fl/fl^). We used an optokinetic tracking system (OptoMotry, Cerebral Mechanics [48]), which evaluates head-tracking responses to moving sine wave gratings projected onto four monitors surrounding the animal. At 4 months of age, mutant animals exhibited no measurable head-tracking behavior across all spatial frequencies tested, indicating a complete loss of visual functions (Fig. 1B).

Because the optomotor response critically depends on the integrity of subcortical visual pathways, and particularly on the function of RGCs projecting to the accessory optic system, we hypothesized that the observed visual impairment might be due to RGC degeneration. To test this, we quantified the total number of RBPMS+ cells in flat-mounted retinas at multiple time points using an automated cell-counting algorithm, previously described by Miesfeld et al. [50], that allowed us to accurately measure all RGCs rather than just those in selected regions.

By postnatal day 30 (P30), there were no significant differences between control and Rax-Cre; Ndufs4^fl/fl^ retinas, with both groups exhibiting approximately 50,000 RGCs/retina, as expected (Fig. 1C-D). However, at 6 weeks of age (P42), we detected a significant reduction in RGC soma numbers (control retinas exhibited 49,024 ± 1,496 RBPMS+ cells vs 32,267 ± 775 in Rax-Cre; Ndufs4^fl/fl^; 34.2% reduction. *P*-value < 0.0001), which progressed to almost a complete loss by 4 months (51,453 ± 2,626 RBPMS+ cells in control mice vs 3,566 ± 1684 in Rax-Cre; Ndufs4^fl/fl^; 93.1% reduction, *P*-value < 0.0001, Fig. 1C-D).

To determine whether the degeneration of RGC somas was accompanied by axonal loss, we performed histological analyses of the retrobulbar optic nerve using paraphenylenediamine (PPD) staining, which selectively labels myelinated axons [54]. Consistent with the RBPMS+ cell counts, there was no significant difference in the overall number of optic nerve axons between control littermates and Rax-Cre; Ndufs4^fl/fl^ animals at P30 (*P*-value= 0.89, Fig. 1E-F). However, by P42, mutant optic nerves showed a marked reduction in axon density (57% reduction, *P*-value < 0.0001), accompanied by clear morphological signs of axonopathy, including swollen or fragmented axons and disrupted myelin sheaths, indicative of an ongoing degeneration (Suppl. Fig 1). By four months (P120), axonal loss was nearly complete, mirroring the near-total loss of RGC somas observed in the retina (94% reduction in axons compared to controls, *P*-value < 0.0001). These findings confirm that mitochondrial dysfunction in RGCs leads to progressive degeneration of both somas and axons, with a similar temporal trajectory.

Although quantification of axonal counts at P30 did not reveal a significant reduction, ultrastructural examination by electron microscopy revealed early abnormalities, including focal axonal swelling and irregularities in the myelin sheath (Suppl. Fig. 1, arrows). These changes were subtle at P30 but became much more pronounced and widespread by P45, suggesting that overt axon loss follows an initial period of structural perturbation and that axons, rather than somas, may represent the earliest site of injury in this model, preceding detectable RGC loss.

### Early degeneration is restricted to the inner retina layers

To further define the spatial and cellular specificity of the retinal degeneration in Rax-Cre; Ndufs4^fl/fl^ mice, we performed histological analyses at P42, the earliest stage of overt RGC degeneration (Fig.2). At this time point, retinal degeneration was primarily restricted to the innermost layers of the retina, with a significant thinning of the inner plexiform layer (IPL, *P*-value = 0.022) and the ganglion cell layer/ retinal nerve fiber layer (GCL/RNFL, *P*-value = 0.0002). No significant changes were observed in the thickness of the other retinal layers, including the outer nuclear layer (ONL), outer plexiform layer (OPL), or inner nuclear layer (INL), when compared to littermate controls, consistent with a selective RGC loss (Fig. 2A-B). Correspondingly, immunohistochemical analyses using markers for specific cell types further confirmed that the other major retinal populations remained largely unchanged (Fig. 2D). Thus, the normal densities of photoreceptors (Otx2+ cells in ONL), cone photoreceptors (Otx2+ Cone Arrestin+ cells), and bipolar cells (Otx2+ cells in INL) were all preserved.

**Figure 2.**
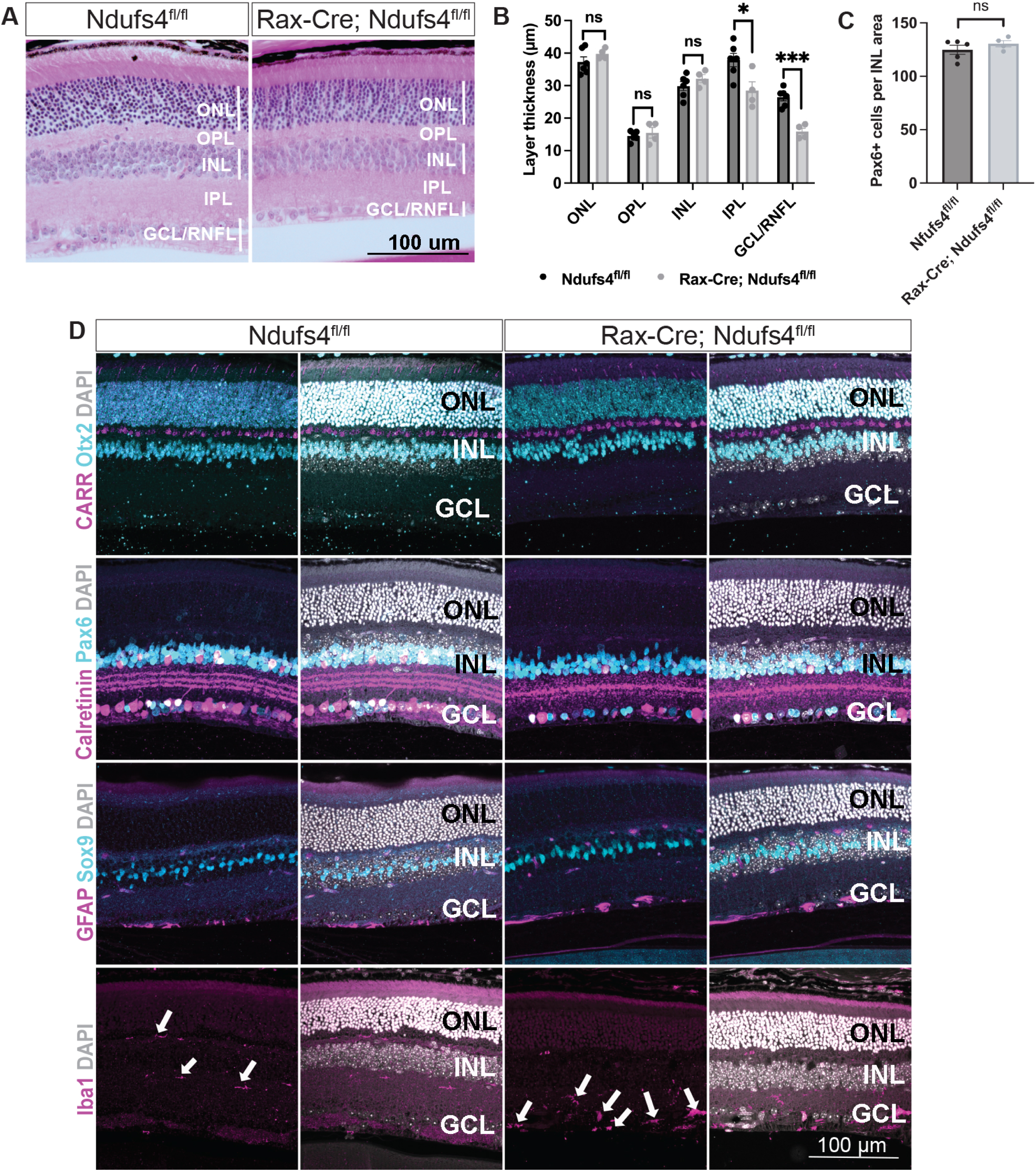
Degeneration in Ndufs4 conditional knockouts was limited to the inner retinal layers. A) H&E stainings of P42 sections. B) Analysis of the thickness of individual retinal layers at P42 (multiple t-tests, n.s. not significant, *P < 0.05, and ***P < 0.001). C) Quantification of Pax6+ cells within the INL, expressed as the number of positive cells per 0.01 mm² of analyzed area (t-test, n.s. not significant). D) Cross-sections of P42 retinas stained with multiple neuronal and immune cell markers reveal alterations in the inner plexiform layer (IPL) and ganglion cell layer (GCL) of Rax-Cre; Ndufs4^fl/fl^ mice, as highlighted by changes in Calretinin and Iba1 stainings. Note the increase in Iba1+ cells in the GCL (arrows).

Previous studies examining whole-body *Ndufs4* knockout mice reported reductions in Calretinin immunolabeling that were attributed to changes in amacrine cell numbers [43, 55]. To evaluate this directly, we quantified amacrine cells by counting INL cells with high Pax6 expression. No significant differences were observed, indicating that amacrine cell numbers remained unchanged in Rax-Cre; Ndufs4^fl/fl^ retinas (*P*-value = 0.33, Fig. 2C-D). However, we detected a significant decrease in Calretinin immunoreactivity, both in somas and within the IPL, consistent with prior observations [43]. In control mice, Calretinin forms a characteristic three-band pattern reflecting the stratified dendritic arborization of specific RGC subtypes and amacrine cells. In Rax-Cre; Ndufs4^fl/fl^ retinas, this stratification was disrupted, with an apparent loss of the distinct stripe organization, and reduced signal intensity in amacrine somas (Fig. 2D). These alterations likely reflect impaired dendritic architecture and synaptic remodeling rather than amacrine cell loss, further supporting the selective impact of mitochondrial dysfunction on RGCs.

To investigate whether these alterations were accompanied by glial responses, we examined Müller glia and microglial morphology and distribution by immunohistochemistry at P42 (Fig. 2D). Surprisingly, despite significant RGC soma and axon loss at this time point, we did not observe obvious signs of Müller glial reactivity. Expression levels of the canonical Müller glia marker Sox9 remained unchanged, and we did not detect any evidence of gliosis, such as hypertrophy or upregulation of GFAP. This lack of reactivity was unexpected, as Müller glia typically respond robustly to retinal stress or injury [56, 57].

In contrast, we observed clear changes in Iba1+ myeloid cell activation and distribution. In control retinas, Iba1+ resident microglia were primarily restricted to the plexiform layers, exhibiting a ramified morphology consistent with a surveillance state. In Rax-Cre; Ndufs4^fl/fl^ mice, however, Iba1+ cells were increased in number and showed an abnormal distribution, particularly within the IPL and GCL/RNFL (arrows, Fig. 2D bottom row). These myeloid cells often displayed a less ramified, more activated morphology, suggesting an early innate immune response at a time point coinciding with the onset of RGC degeneration.

### Accumulation and activation of Iba1+ myeloid cells precede RGC loss

To determine whether the accumulation of Iba1+ myeloid cells observed is a secondary response to an ongoing RGC degeneration or a contributing factor in the disease progression, we examined retinal flat-mounts at P30, prior to visible RGC soma or axonal loss. We quantified myeloid cells using two complementary measures: nearest neighbor distance, which captures spatial clustering and thus reflects localized activation, and the number of Iba1⁺ cells per mm², which reflects overall cell density. Both analyses revealed a significant increase in Iba1+ cells in both the IPL and GCL/RNFL in Rax-Cre; Ndufs4^fl/fl^ retinas compared to littermate controls (Fig. 3A-E). This accumulation was evident across the entire retina, including both peripheral and central regions, indicating a widespread early immune response. In addition to greater numbers and clustering, myeloid cells displayed activated morphologies, particularly in the central retina surrounding the ONH, where we observed rod-shaped, elongated cells in close proximity to RGC axon bundles (Fig. 3F).

**Figure 3.**
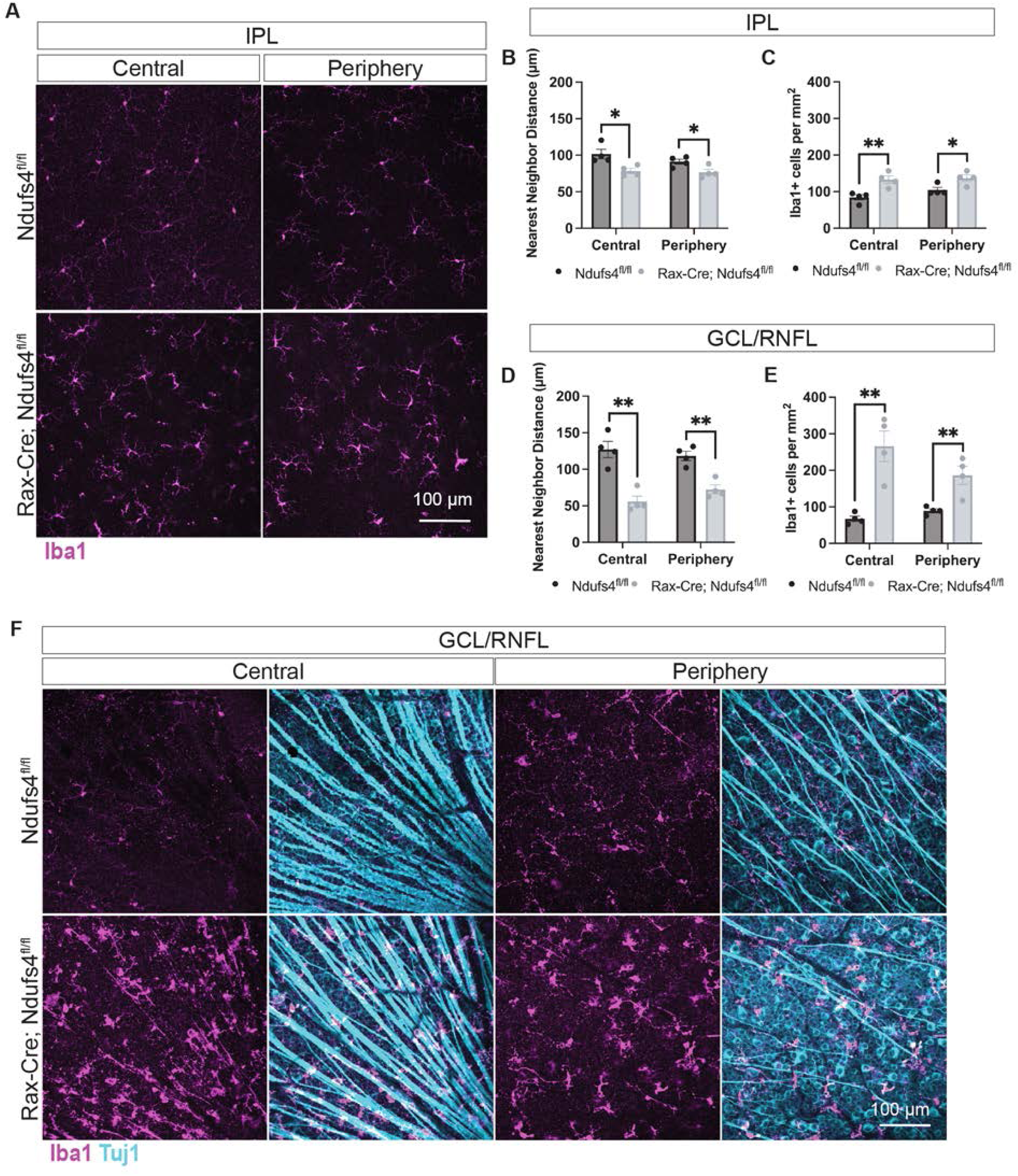
Rax-Cre; Ndufs4^fl/fl^ mice showed an increase in myeloid cells by P30. A) Stanings of retinal flatmounts show an increase of myeloid cells (Iba1) in both the center and periphery of the IPL of Rax-Cre; Ndufs4^fl/fl^ mice. B to E) Quantification of nearest neighbor distance and number of Iba1+ cells per mm^2^ in the IPL and GCL/RNFL of Iba1-stained retinal flat-mounts. Central and peripheral retina were quantified separately for each layer (multiple t-tests, *P < 0.05 and **P < 0.01). F) Flat-mount stainings showed an increase of myeloid cells (Iba1, magenta) in both the center and periphery of the GCL/RNFL of Rax-Cre; Ndufs4^fl/fl^ mice. Morphologies characteristic of reactive myeloid cells were observed near RGC axons (Tuj1, teal).

Since Iba1 is a general marker for myeloid lineage cells, these analyses did not distinguish resident microglia from infiltrating macrophages. However, the early and diffuse increase observed suggests that immune cell activation occurs before overt RGC degeneration and may play a role in the pathological process rather than being merely a consequence.

### Myeloid cell depletion protects RGCs from degeneration

To directly test whether the myeloid cell response observed contributes to RGC degeneration, we pharmacologically depleted myeloid lineage cells using PLX5622, a well-established CSF1R inhibitor [58]. The treatment was initiated at weaning age (P21), before the onset of detectable neurodegeneration, by administering a chow formulated with PLX5622 (Research Diets, Inc). For each animal, effective myeloid depletion was confirmed by immunohistochemical analysis of Iba1+ cells in retinal flat-mounts (Fig. 4A). RGC survival was then assessed in the other eye and compared to Rax-Cre; Ndufs4^fl/fl^ littermates fed the same base diet (AIN-76A) but lacking the drug. Interestingly, PLX5622-treated animals exhibited a significant and widespread neuroprotective effect, with substantially higher numbers of surviving RGCs at P45 relative to untreated mice (41,572 ± 2,370 RBPMS+ cells in retinas treated with PLX5622 vs 28,355 ± 2,958 RBPMS+ in untreated mice, *p*-value = 0.0014, Fig. 4B-D). These findings demonstrate that activation of the innate immune system is not merely a secondary response to RGC loss, but rather a required contributor to the pathological mechanism driving neurodegeneration in this model.

**Figure 4.**
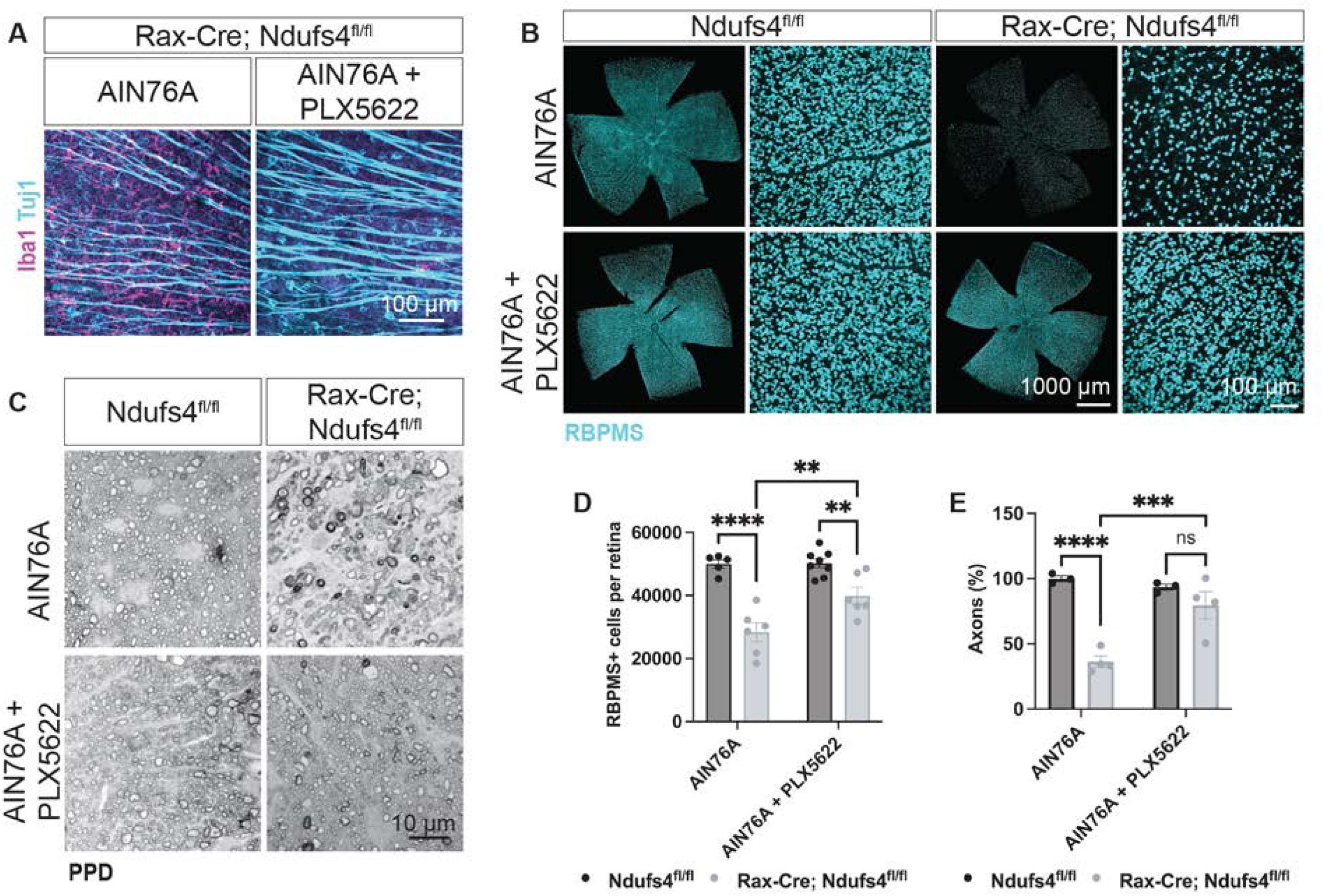
Depletion of myeloid cells in Rax-Cre; Ndufs4^fl/fl^ mice improves RGC cell survival. A) P45 retinal flatmounts stained with Iba1 (myeloid cells, magenta) and Tuj1 (RGCs, teal), showed near total ablation of myeloid cells in the GCL/RNFL of Rax-Cre; Ndufs4^fl/fl^ mice fed with PLX5622. B) Retinal flat-mounts stained with RBPMS (RGCs, teal). Note the rescue of the RGC degeneration in Rax-Cre; Ndufs4^fl/fl^ mice upon PLX5622 treatment. C) Paraphenylenediamine (PPD) staining in retrobulbar optic nerve sections showed axonal rescue in Rax-Cre; Ndufs4^fl/fl^ mice fed with PLX5622. D and E) Quantification of RBPMS+ cells per retina and axon percentages in animals depleted of myeloid cells with PLX5622 treatment (two-way ANOVA, n.s. not significant, **P < 0.01, ***P < 0.001, and ****P < 0.0001).

We also assessed the impact of PLX5622 treatment on optic nerve integrity using PPD staining, as described above. Consistent with our RBPMS counts of RGC somas, depletion of myeloid cells provided a strong protection of RGC axons in Rax-Cre; Ndufs4^fl/fl^ mice (Fig. 4C-E). In untreated samples, axonal counts were reduced by 63% compared to littermate controls, whereas PLX5622-treated Rax-Cre; Ndufs4^fl/fl^ showed only a 14% reduction (*P-*value = 0.0007, untreated vs. treated).

Overall, these findings indicate that myeloid cell activation contributes directly to RGC degeneration in Ndufs4 deficiency, and that targeting this response can mitigate both soma and axon loss.

### Mosaic Ndufs4 deletion reveals cell-autonomous vulnerability of RGCs

Our findings from PLX5622-treated animals demonstrated that myeloid cell depletion confers robust neuroprotection, suggesting that innate immune activation is necessary for RGC loss. However, it remains unclear whether myeloid cells selectively eliminate compromised RGCs or whether they also contribute to the death of otherwise healthy neurons through non-cell-autonomous mechanisms. To address this question, we employed a genetic strategy to cross Ndufs4^fl/fl^ mice with αPax6-Cre, a mosaic Cre-driver expressed under the α-enhancer element of the Pax6 [47]. In this model, Cre activity is restricted to the peripheral retina from early stages of development, enabling spatially defined deletion of *Ndufs4* and allowing for direct comparison of mutant (Cre+, peripheral) and wild-type (Cre-, central) regions within the same retina (Fig. 5A) Behaviorally, αPax6-Cre; Ndufs4^fl/fl^ animals (hereafter α-Cre; Ndufs4^fl/fl^) exhibited a measurable reduction in visual acuity by four months of age as assessed by optomotor responses, but the degree of visual loss was milder than the complete loss observed in the pan-retinal Rax-Cre; Ndufs4^fl/fl^ model, consistent with partial mosaic recombination (21.4% reduction in visual acuity, *P*-value = 0.0017, Fig. 5B).

**Figure 5.**
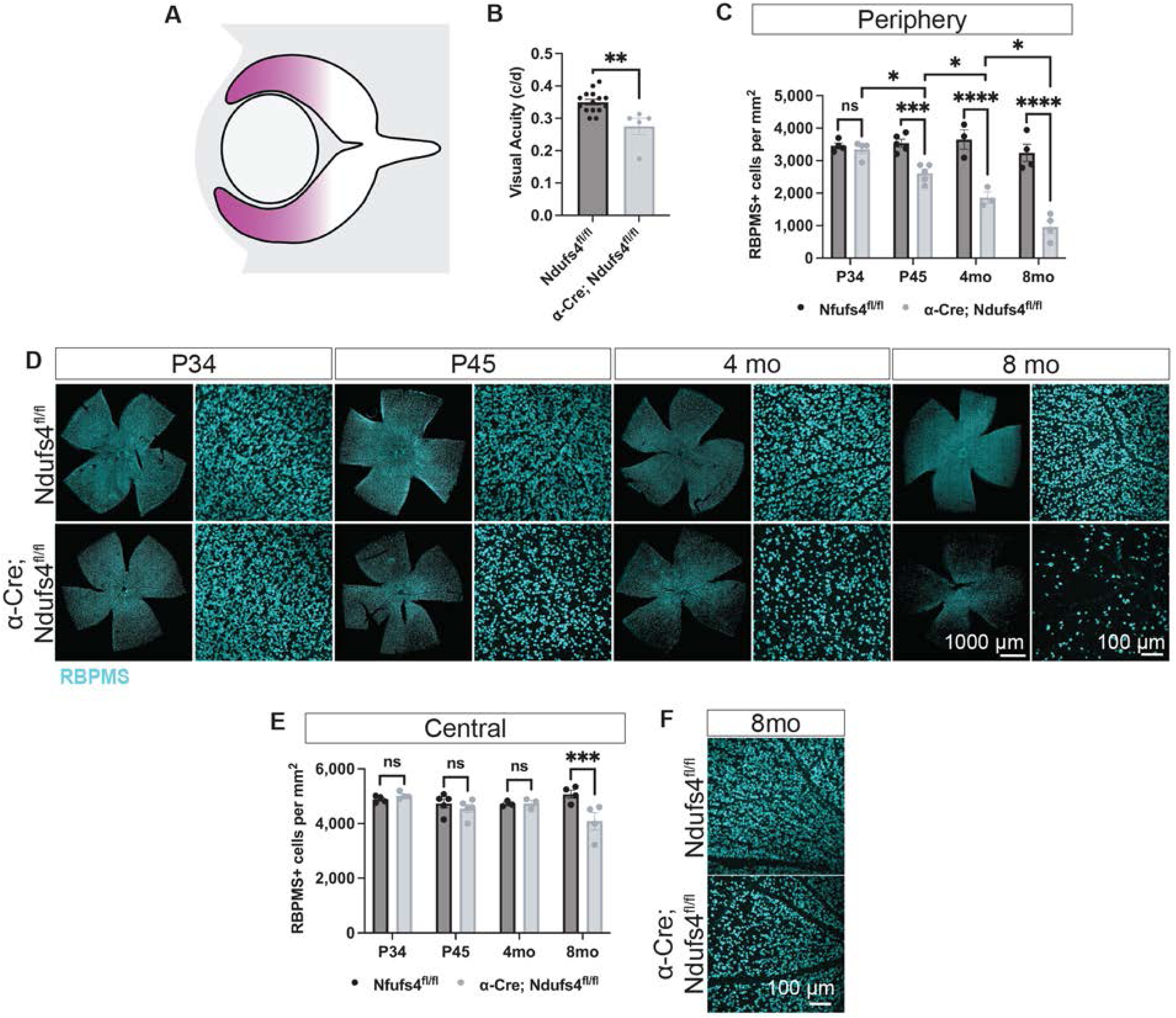
RGC degeneration is limited to peripheral α-Cre; Ndufs4^fl/fl^ retinas at early time-points. A) Simplified diagram of the embryonic retina showing the expression pattern of Cre (magenta) in α-Cre mice. B) Optomotor response of 4-month animals (t-test, **P < 0.01). C) Quantification of RBPMS+ RGCs per mm^2^ in the peripheral retina over time (two-way ANOVA, n.s. not significant, *P < 0.05, ***P < 0.001, and ****P < 0.0001). D) Flat-mounts labeled with RBPMS (teal) showed progressive RGC loss in the periphery of α-Cre; Ndufs4^fl/fl^ retinas. Close-up panels represent the peripheral retina. E) Quantification of RBPMS+ RGCs per mm^2^ in the central retina over time (two-way ANOVA, n.s. not significant and ***P < 0.001). F) RBPMS staining of flat-mounted retinas showed loss of RGCs in the central retina of α-Cre; Ndufs4^fl/fl^ animals by 8 months.

Histological analyses revealed that degeneration first emerged by P45, consistent with our previous results, and was spatially confined to peripheral Cre+ regions, where *Ndufs4* is deleted. Thus, we did not observe any changes by P30 (3,454 ± 89 and 4,886 ± 63 RBPMS+ cells/mm^2^ in the periphery and central areas of control retinas, respectively vs 3,347 ± 135 and 5,020 ± 62 RBPMS+ cells/mm^2^ in α-Cre; Ndufs4^fl/fl^, *P*-values = 0.67 and 0.57); by P45, we observed a 26% reduction in RGC counts in the Cre+ regions (3,540 ± 120 RBPMS+ cells/mm^2^ in the periphery of control retinas *vs* 2,608 ± 126 in α-Cre; Ndufs4^fl/fl^, *P*-value < 0.0001). In contrast, RGCs in the central Cre-areas remained structurally intact at this stage, despite exposure to the same microenvironment (Fig. 5C-D, 4,737 ± 161 RGCs/mm^2^ in controls *vs* 4,550 ± 148 in α-Cre; Ndufs4^fl/fl^, *P*-value = 0.38). By 4 months, we observed the same effect where the degeneration was restricted to the Cre+ regions but the degeneration had progressed further (from a 26% reduction at P45 to a 49% reduction at P120, *P*-value < 0.0001). Interestingly, at later stages, RGC degeneration extended into the central Cre-regions of the retina, indicating that disease propagation can occur beyond the initial zone of genetic insult (Fig. 5E-F). Thus, at 8 months, the central retina exhibited 5,077 ± 146 RGCs/ mm^2^ in controls vs 4,085 in α-Cre; Ndufs4^fl/fl^, representing a 19.5% loss, *P*-value = 0.0003). This delayed degeneration in genetically intact areas strongly supports the presence of a secondary, non-cell-autonomous bystander effect, likely driven by inflammatory mechanisms initiated in Cre+ regions.

We also examined optic nerves from α-Cre; Ndufs4^fl/fl^ mice at four months of age using PPD staining to assess axonal integrity. Consistent with the mosaic retinal phenotype, overall axonal loss in these animals was less severe than in Rax-Cre; Ndufs4^fl/fl^ mice, but degeneration was nonetheless evident (Suppl. Fig. 2). Notably, axonal loss was observed in both the central and peripheral regions of the nerve, with a more substantial reduction in axon density in the peripheral portion (48% reduction in the periphery, *P* = 0.0009 and 26% axonal loss in the central part of the nerve, *P* = 0.006). While the spatial organization of axons within the optic nerve suggests that degeneration is more pronounced among fibers originating from peripheral, Cre+ RGCs, it does not exclude the possibility that Cre-RGCs eventually contribute to axonal loss, particularly as degeneration progresses. Overall, these findings support a model in which cell-intrinsic mitochondrial dysfunction initiates axonal degeneration, which later spreads to affect adjacent, genetically intact neurons.

### Innate immune responses in the mosaic **α**-Cre; Ndufs4^fl/fl^ model were detected in both the central and peripheral retina

To further investigate the spatial and temporal dynamics of innate immune activation in the mosaic α-Cre; Ndufs4^fl/fl^ model, we analyzed the distribution and organization of Iba1+ myeloid cells at P30 and P45. At P30, prior to detectable RGC soma loss, we observed early changes in Iba+ cells (Fig. 6A-E). At this stage, increased Iba1 signal and clustering were restricted to the GCL/RNFL, with no significant changes in the IPL. However, in the periphery, we did detect a reduction in nearest neighbor distance among Iba1+ cells in the IPL, suggesting localized myeloid cell activation may begin earlier in Cre+ zones. Despite these subtle regional differences, innate immune activation was observed in both Cre+ and Cre-regions.

**Figure 6.**
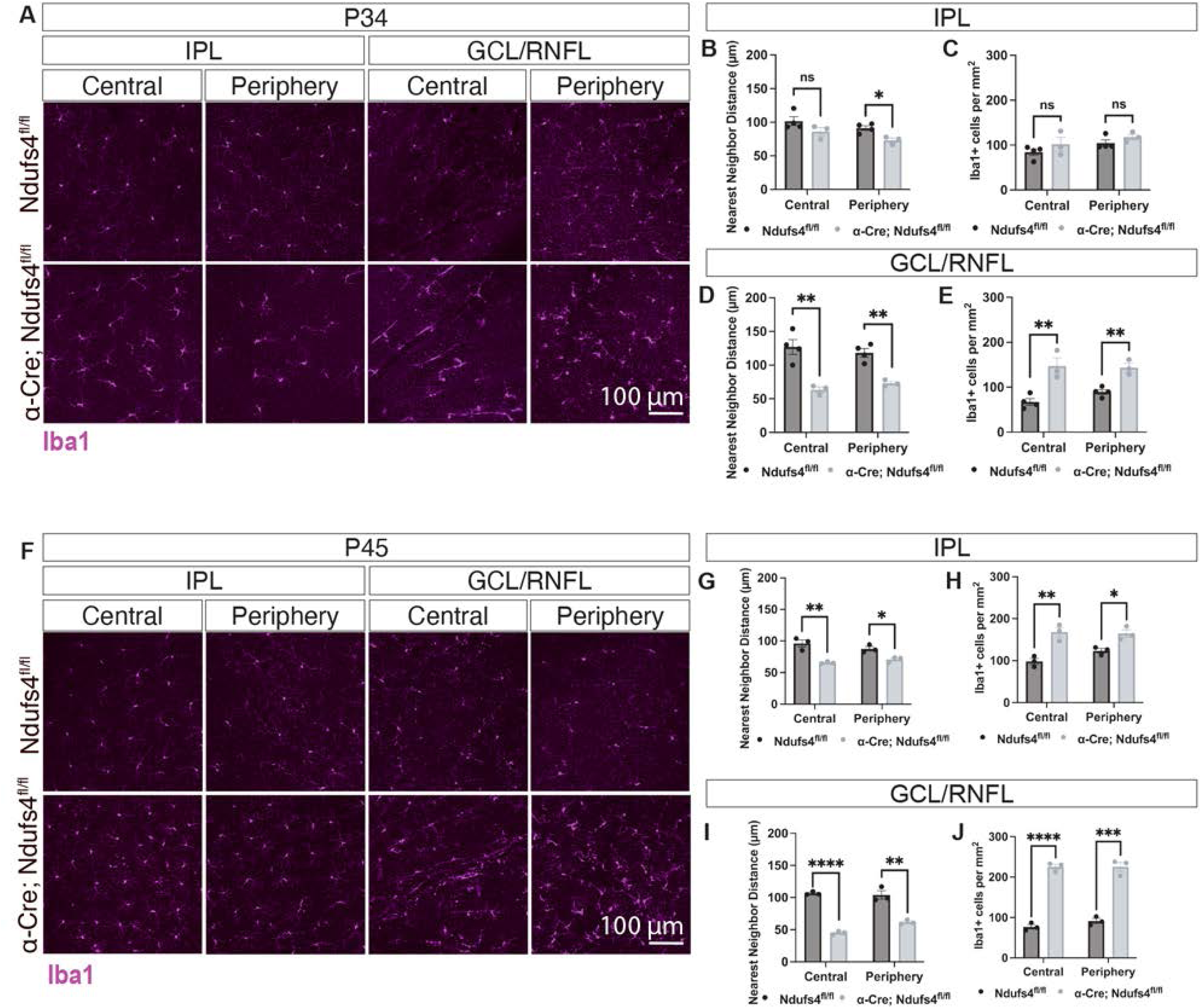
Myeloid cell populations are altered in both peripheral and central retina of α-Cre; Ndufs4^fl/fl^ mice. A) α-Cre; Ndufs4^fl/fl^ retinal flat-mounts show changes in the morphology and distribution of Iba1+ cells (magenta) in both IPL and GCL/RNFL of P30 retinas. Note that changes are not limited to the peripheral retina but also affect the central regions. B to E) Quantification of nearest neighbor distance and number of Iba1+ cells per mm^2^ in the IPL and GCL/RNFL of Iba1-stained P34 retinal flat-mounts (multiple t-tests, n.s. not significant, *P < 0.05, and **P < 0.01). F) α-Cre; Ndufs4^fl/fl^ retinal flat-mounts stained with Iba1 (magenta) show an increase in the number of myeloid cells by P45; these changes are widely distributed throughout the IPL and GCL/RNFL. G to J) Quantification of nearest neighbor distance and number of Iba1+ cells per mm^2^ in the IPL and GCL/RNFL of Iba1-stained retinal flat-mounts at P45 (multiple t-tests, *P < 0.05, **P < 0.01, ***P < 0.001, and ****P < 0.0001).

At P45, we detected a robust increase in the number of Iba1+ cells in both the central and peripheral retina, despite the degeneration remained confined to Cre+ peripheral regions. This increase was observed across multiple retinal layers, including the IPL and the GCL/RNFL (Fig. 6F-J). Similarly, we examined the expression of CD68, a lysosomal membrane protein commonly upregulated in phagocytic and activated myeloid cells. By P45, CD68 immunoreactivity was markedly elevated in nearly all Iba1+ cells, consistent with a shift from a homeostatic to an activated or reactive state (Suppl. Fig. 3).

These findings indicate that immune activation arises before cell-intrinsic degeneration and extends beyond it, implicating the innate immune system as an active driver of disease progression. In this model, inflammation emerges early and spreads broadly, without strictly following the pattern of RGC loss or remaining limited to genetically targeted (Cre+) regions.

To assess the functional role of myeloid cells in mediating RGC degeneration in the α-Cre; Ndufs4^fl/fl^ model, we treated animals with PLX5622 starting at weaning (P21) to deplete myeloid populations, as described above. Similar to our observations in the Rax-Cre model, PLX5622 treatment resulted in a significant neuroprotective effect, with greater preservation of RGCs compared to control animals receiving the same diet without the drug (Suppl. Fig. 4). This finding confirms that innate immune cells contribute to RGC loss in this mosaic model, further supporting the hypothesis that immune activation is a key pathological driver rather than merely a consequence of degeneration.

### Inflammation decreases at later stages despite ongoing and expanding RGC degeneration

We next examined inflammatory changes at later disease stages, motivated by the observation that RGC loss not only progresses beyond P120 but also expands into regions that retain Ndufs4 expression (Fig 5). In α-Cre; Ndufs4^fl/fl^ mice, the number of Iba1+ myeloid cells was markedly reduced by 4 months of age compared to earlier time points. Although inflammation was not fully resolved at this stage (Fig. 7), the overall myeloid response was substantially diminished relative to the early phases of degeneration. For instance, at P45, the number of Iba1+ cells in the central retina increased 2.93-fold in α-Cre; Ndufs4^fl/fl^ mice (from 76 ± 9 Iba1+ cells in the GCL in controls to 225 ± 11 cells in α-Cre; Ndufs4^fl/fl^, Fig 6F, J), while at later stages this difference was 1.3-fold (from 81 ± 9 Iba1+ cells in the GCL in controls to 108 ± 13 cells in α-Cre; Ndufs4^fl/fl^, Fig 7A, E).

**Figure 7.**
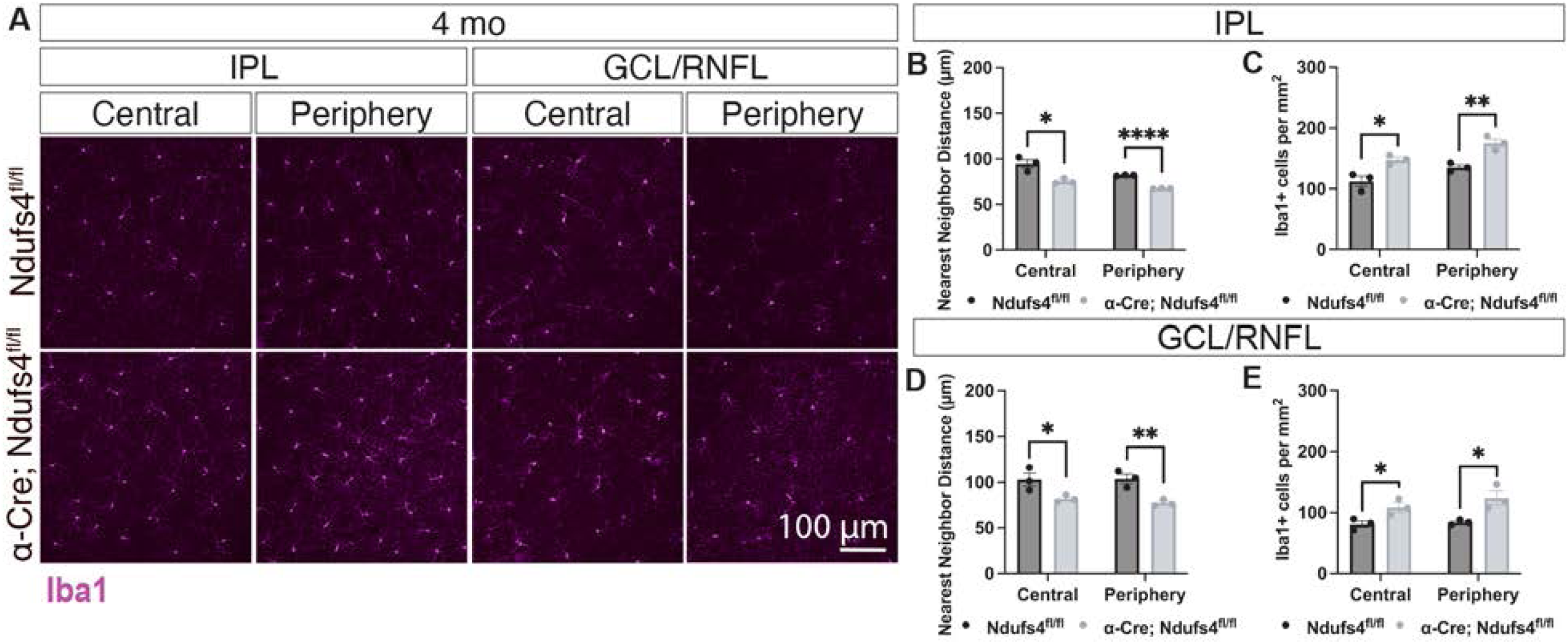
Inflammation decreases by 4 months in α-Cre; Ndufs4^fl/fl^ retinas. A) Iba1 stainings of α-Cre; Ndufs4^fl/fl^ retinal flat-mounts indicate that myeloid cell numbers and distribution are close to control levels by 4 months of age. B to E) Quantification of nearest neighbor distance and number of Iba1+ cells per mm^2^ in the IPL and GCL/RNFL of Iba1-stained retinal flat-mounts at 4 months (multiple t-tests, *P < 0.05, **P < 0.01, and ****P < 0.0001).

Notably, despite this reduction in Iba1+ cell density and clustering, RGC degeneration continued to progress between 4 and 8 months and spread into Cre-regions of the retina (Fig. 5). These findings suggest that the early inflammatory response is not sustained at later stages, yet RGC degeneration persists and expands, indicating that additional or secondary mechanisms contribute to disease progression beyond the initial inflammatory phase.

### Temporal shift from chemokine- to complement-associated inflammation during the progression of RGC pathology

To investigate stage-specific inflammatory changes during RGC degeneration, we collected retinas at P30 and P120 and analyzed their cytokine proteomic profiles using a cytokine array kit (R&D, Fig. 8). P30 was selected to capture early inflammatory events, before substantial RGC death had occurred. P120 was chosen to examine the later stages, when RGC degeneration extends into Cre-regions, allowing us to assess the inflammation profiles associated with disease progression.

**Figure 8.**
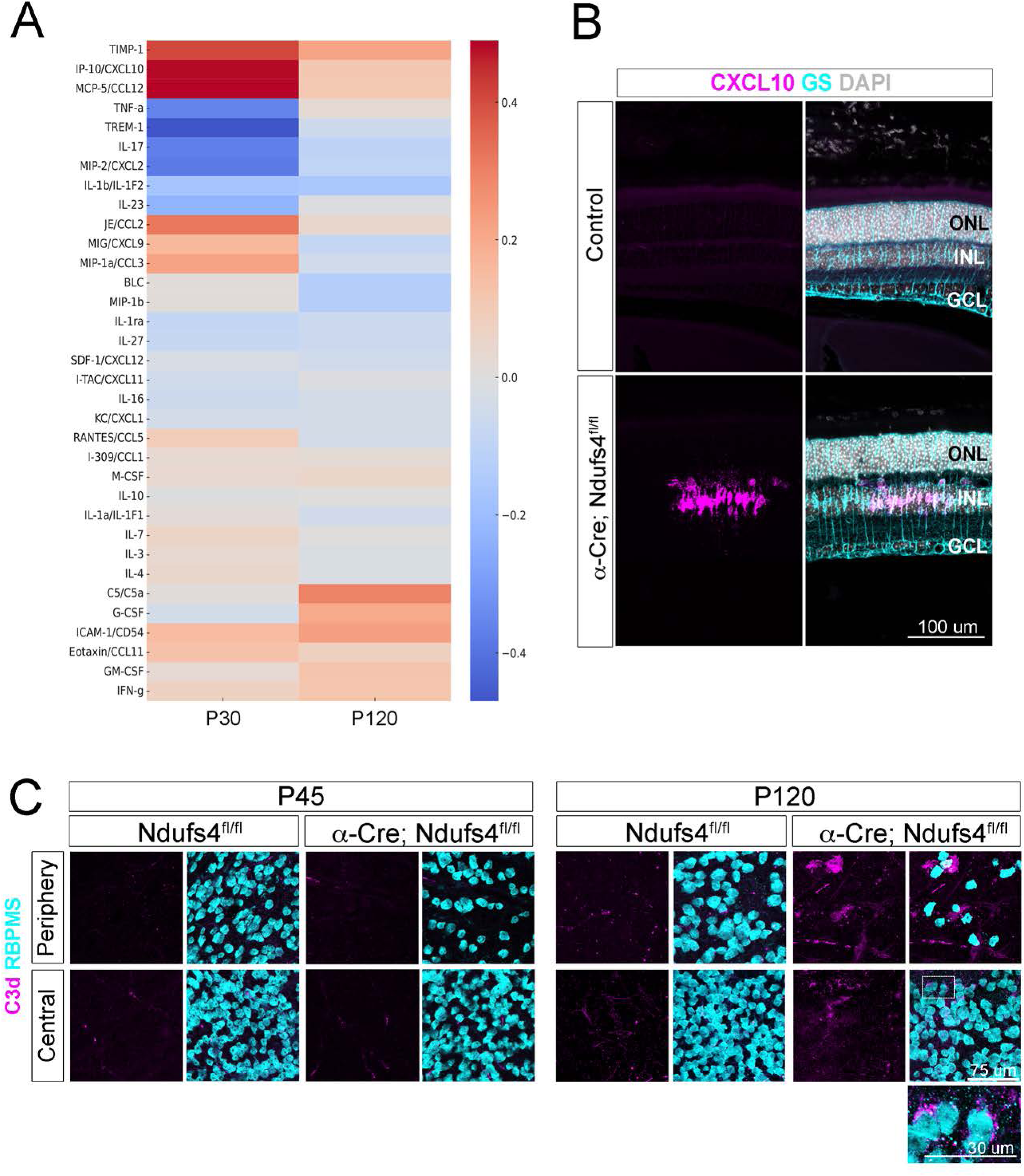
Cytokine proteome profiling identifies a switch from early to late stages of RGC pathology. **A)** Hierarchical clustering heatmap of cytokine expression changes compared to Ndufs4^fl/fl^ controls at P30 and P120. Each data point represents the average of two independent experiments, each conducted with two technical duplicates. B) CXCL10 *in situ* hybridization (magenta) was performed on P45 retinal sections and the samples were then co-stained with RBPMS (teal) and DAPI (gray). C) RBPMS (teal) and the complement protein C3 (magenta) staining on flat-mounted retinas at P45 and P120. Note the accumulation of C3 in α-Cre; Ndufs4^fl/fl^, particularly around RGCs as highlighted in the close-up panel (bottom right).

At early stages, cytokine profiling revealed strong upregulation of CXCL10, CCL12, CCL2, CCL3, CXCL9, and CCL11, together with high levels of TIMP-1 and ICAM-1 (Fig. 8A). To validate these findings, we analyzed the expression of CXCL10, one of the cytokines with largest early changes in expression, by *in situ* hybridization (Fig. 8B). Consistent with the array data, we did not observe any signal in controls, whereas α-Cre; Ndufs4^fl/fl^ samples contained discrete patches of Müller glia with pronounced CXCL10 expression.

Overall, this chemokine milieu is consistent with a pro-inflammatory environment capable of supporting both activation of resident myeloid cells and recruitment of additional immune populations, including monocytes/macrophages and CXCR3+ lymphocytes. In this direction, the changes in cytokine expression coincided with a widespread increase in Iba1+ cells (Fig. 6). However, to further assess whether lymphocytes were also recruited to the diseased retina, we stained flat-mounted retinas with CD3, which labels all T cells. CD3+ cells were readily observed throughout the retina, demonstrating that T lymphocytes are recruited in this model (Suppl. Fig.5). By late stages of disease progression (P120), most of these chemokines (CCL12, CCL2, CCL3, CXCL9, CXCL10, and CCL11) declined toward baseline levels and the inflammatory profile shifted toward complement activation, reflected in elevated C5a, along with increased G-CSF (Fig. 8A). CD3 staining revealed that T cells remained present at this stage (Suppl. Fig. 5), while complement component C3d became detectable in patches and surrounding RGCs, a feature not observed at earlier stages (Fig.8C).

This pattern pointed to a shift from an early chemokine-driven phase that recruits and activates immune cells to a later effector phase, which includes complement activation, potentially perpetuating retinal injury even after local myeloid cell numbers return toward baseline. Together, these findings provide a stage-specific molecular signature of the evolving inflammatory response and indicate that the processes that drive RGC degeneration are not static but evolve from an early recruitment phase to a later injury-amplifying phase.

## Discussion

We have generated two complementary models of *Ndufs4* deficiency that capture the progression of mitochondrial optic neuropathy: one exhibiting pan-retinal involvement and the other showing a sectorial, mosaic pattern. Unlike previous *Ndufs4* models, which are limited by very short lifespans [40, 43, 45], these novel systems enable detailed analyses of the temporal dynamics of RGC degeneration and the relative contributions of cell-autonomous and non-cell-autonomous mechanisms.

Notably, our transgenic models revealed a phased progression of retinal neurodegeneration, which we have categorized into three distinct stages: early inflammatory responses precede detectable neuronal loss, followed by cell-autonomous degeneration of RGC somas and axons, and finally a non-cell-autonomous spread of pathology. This sequential framework provides a unique platform to analyze the cellular and molecular mechanisms that drive disease at each stage.

### First Phase: Pre-Degeneration stage

In the earliest phase (around P30), the first detectable abnormalities appeared in the optic nerve, where we observed sporadic myelin irregularities and axonal swelling. These findings suggest that axons, rather than somas, are the initial site of injury in this model. Axons are highly dependent on mitochondrial transport to meet local energy demands. Recent data using genetic and optic nerve injury models have identified failure of mitochondrial transport as a central event in RGC pathology [10, 59, 60]. In this direction, loss of *Ndufs4* could compromise axonal ATP production and calcium buffering earlier and more severely than in somas, thereby impairing axoplasmic transport, disrupting myelin-axon interactions, and rendering axons vulnerable to structural stress. One possible consequence of such axonal dysfunction is the local release of damage-associated molecular patterns (DAMPs), such as extracellular ATP or oxidized mtDNA that, in turn, are known to rapidly mobilize and activate innate immune sensors [61, 62]. Consistent with this idea, at early stages of the pathology, before any measurable loss of RGC somas or axons, we detect a robust increase in Iba1+ myeloid cells across both Cre+ and Cre-retinal regions.

Also at these early stages of the pathology, we detect an upregulation of cytokines and chemokines, including ICAM-1, CXCL10, CXCL9, CCL2, CCL3, CCL12, CCL5, and TIMP-1. These molecules are known to promote the adhesion, trafficking, and activation of monocytes and leukocytes [63–67]. Notably, we validated CXCL10 (also known as IP-10) expression by *in situ* hybridization and identified Müller glia as a source, despite the absence of detectable Müller glia reactivity (GFAP). The patchy nature of CXCL10 expression indicates heterogeneous responses within the Müller glia population, with some but not all cells expressing this chemokine. In particular, CXCL10 is upregulated in models of retinal injury or increased intraocular pressure where it signals via CXCR3 to recruit inflammatory cells (both macrophages and leukocytes), contributes to ER stress, and RGC loss [68–71]. Similarly, CCL2 (JE) and CCL12 (MCP-5) are well-known drivers of monocyte recruitment [71–74], ICAM-1 (CD45) facilitates leukocyte diapedesis [75], and TIMP-1 may reflect early extracellular matrix remodeling in response to axonal stress [76]. Collectively, these molecular cues suggest that early inflammatory signaling is not only increasing immune cell number but may also be shaping their behavior and phenotype. A previous study indicated that CXCL10 depletion does not significantly affect the onset or progression of neurologic symptoms and weight loss in Ndufs4^-/-^ mice [77], but the effects on retinal degeneration have not been investigated. Similarly, using the *Ndufs4* full knockout model, a previous study has shown that the disruption of the adaptive immune system does not affect disease progression in the brain [78].

Around the ONH, Iba1+ cells exhibit distinct morphologies characterized by elongated cell bodies and extended processes that align along nerve fibers. Similar rod-like morphologies have previously been associated with transcriptional states resembling disease-associated microglia (DAMs), suggesting that these structural changes could coincide with a shift toward neurodegeneration-linked phenotypes [79–81]. A rod-shaped microglial population has been previously shown to participate in debris clearance following axonal injury [82, 83]; thus, some of the cells at the retinal center could be engulfing Cre+ axonal material originating from peripheral RGCs.

Taken together, focal axonal stress may initiate a paracrine inflammatory cascade that engages resident and/or infiltrating myeloid cells, and also recruits the adaptive immune system, though the specific contribution of the latter to RGC degeneration has yet to be determined.

### Second phase: degeneration of axons and cell bodies in vulnerable areas

The second phase (P45–120) is characterized by a sharp decline in RGC somas and axons, which is restricted to Cre+ regions, consistent with cell-autonomous vulnerability. This period coincides with peak numbers of Iba1+ myeloid cells, which also show high CD68 expression, reflecting phagocytic activity.

Myeloid cells at this stage contribute directly to RGC death, possibly through the release of proinflammatory cytokines, oxidative stress, or the phagocytosis of stressed but potentially viable neurons. The protective effect of PLX5622-mediated depletion of myeloid cells provides strong evidence that this population plays a causal role, supporting the idea that myeloid activity is a key amplifier of the cell-autonomous mitochondrial defect. Consistent with these findings, inhibition of CSF1R has been previously shown to extend survival as well as rescue several symptoms such as seizures, motor deficits, and breathing alterations in Ndufs4 knockout models [84, 85].

These findings support a two-hit model in which the initial degeneration occurs only when intrinsic mitochondrial defects coincide with myeloid-mediated inflammatory activity, whereas either factor alone is insufficient to drive widespread cell loss.

### Third phase: secondary, non-cell-autonomous damage

In the third phase (> 4 months), RGC degeneration spreads into Cre- regions, despite a decline in Iba1+ cell numbers to near-control levels within the retina. This dissociation between myeloid presence and ongoing degeneration suggests that once sufficient damage and inflammatory signaling accumulate, additional mechanisms sustain the spread of pathology. At this stage, proteomic cytokine analysis reveals persistent elevation of ICAM-1, along with a late rise in complement component C5a. The complement is a potent mediator of immune recruitment and local tissue injury and can promote degeneration through complement opsonization, membrane attack complex (MAC) formation, or sustained low-level inflammatory signaling. These findings raise the possibility that complement activation may participate in disease progression once initial axonal injury and myeloid activation have set the stage. In line with this, previous work has demonstrated that complement activation contributes to RGC injury. Human studies showed increased expression of complement components in the glaucomatous retina [86, 87]. C1q, produced primarily by microglia, initiates the classical complement cascade, and complement activation often occurs near stressed RGCs and their synapses, promoting aberrant pruning, opsonization, and bystander degeneration [88, 89]. In agreement with this, C1q deficiency is neuroprotective [20], and inhibition of C3 activation using an AAV2 therapeutic approach significantly delayed disease progression in the DBA/2J glaucoma model [90]. Our findings are consistent with a model in which complement could serve as a mechanistic bridge between the early cytokine-driven stress response and the later, non-cell-autonomous neuronal damage, though this has not yet been directly examined.

Notably, some of the sequence of events detailed here parallel observations in human pathology. In LHON, mitochondrial dysfunction preferentially affects small-caliber axons of the optic nerve, with axonal swelling and degeneration preceding RGC soma loss [91]. Similarly, in glaucoma models, structural and functional changes in axons at the optic nerve head are among the earliest detectable abnormalities, with soma degeneration occurring later [92–94]. These comparisons suggest that axonal vulnerability may represent a common initiating event in optic neuropathies of diverse etiology, particularly when mitochondrial dysfunction is involved. Our data extend this concept by demonstrating how early mitochondrial stress can trigger a cascade of myeloid activation and, ultimately, cell-autonomous propagation of degeneration.

Taken together, these results define a sequence of events and reveals two different mechanisms of cell death. Importantly, this phased model provides a framework for considering therapeutic timing. In the prodromal stage, when axonal stress is detectable, but cell loss has not yet occurred, interventions aimed at stabilizing mitochondrial function or improving axonal bioenergetics may hold the most significant potential. During the active degeneration phase, our findings indicate that pharmacological depletion of myeloid cells with PLX5622 provides strong neuroprotection establishing that this population is a key driver of disease progression. While PLX5622 itself is not a viable therapeutic, this proof-of-principle result highlights the importance of myeloid biology and further suggests that strategies aimed at modulating myeloid activity could be beneficial if targeted appropriately. Finally, at later stages, the emergence of complement activation raises the possibility that complement pathways contribute to the spread of degeneration.

The data presented revealed a stepwise mechanism of mitochondrial optic neuropathy, providing a coherent framework for understanding both the cellular origins and the evolving mechanisms of neurodegeneration. Extending these observations, future single-cell RNA sequencing will decipher the full spectrum of immune and glial responses across disease stages, while functional studies will be needed to determine the specific contributions of individual cytokines, the adaptive immune system, and complement components to RGC injury and degeneration.

## Conclusion

This study demonstrates that mitochondrial optic neuropathy progresses through a phased sequence, in which early axonal and retinal stress coincides with the activation of myeloid cells, followed by the degeneration of RGC somas and axons, and ultimately, a spread of pathology. These findings highlight the interplay between intrinsic mitochondrial dysfunction and non-cell-autonomous inflammatory mechanisms, providing a framework for understanding the cellular origins and progression of optic neuropathies such as LHON. Importantly, our models define distinct stages that could be therapeutically targeted by supporting mitochondrial function and limiting inflammation early, modulating myeloid activity during active degeneration, and addressing complement-mediated propagation at later stages, offering concrete insights for the development of temporally informed interventions.

## Abbreviations

DOA: Dominant optic atrophy
GCL: Ganglion cell layer
INL: Inner nuclear layer
IPL: Inner plexiform layer
LHON: Leber’s hereditary optic neuropathy
ONL: Outer nuclear layer
OPL: Outer plexiform layer
P: Postnatal day
RGC: Retinal Ganglion Cell
RNFL: Retinal nerve fiber layer
RPE: Retinal pigment epithelium

## Acknowledgments

We thank Keiko Hino for technical assistance and all members of our laboratories for their helpful insights. We also thank Drs. Nadean Brown and Tom Glaser for valuable comments and generosity with reagents. This study was supported by The Glaucoma Research Foundation Catalyst for a Cure grant to ALT, XD, YH and DSW, the UCD Team Science Award to ALT and NMA, 1R21EY037445 to NMA, and NIHR01EY030138 to XD. ES is Ramon y Cajal fellow (RyC2019-028501-I) and received funds from MICIU Proyectos I+D+i “Retos Investigacion” (RTI2018-101838-J-I00) and Ministerio de Ciencia e Innovación (MICINN) (PID2019-107633RB-I00 and PID2022-142544OB-I00). AQ received funds from MICIU PID2020-114977RB-I00, PID2023-151947OB-I00, AGAUR (2017SGR-323, 2021SGR-720), and “la Caixa” Foundation (ID 100010434), under the agreement 1183 LCF/PR/HR20/52400018. AQ is a recipient of an ICREA Academia award. We benefited from the use of the National Eye Institute Core Facility for histology sample processing and EM experiments [supported by P30 EY012576].

## Supplemental Figure legends

**Suppl. Figure 1. Ndufs4 loss results in defects of the myelin sheath.** Electron microscopy images of retrobulbar optic nerves show disrupted myelin sheaths (arrows) in both P30 and P45 Rax-Cre; Ndufs4^fl/fl^ animals.

**Suppl. Figure 2.** α**-Cre; Ndufs4^fl/fl^ animals show axonal degeneration at 4 months.** A) Paraphenylenediamine (PPD) staining to label myelinated axons in optic nerve sections demonstrates axonal loss in α-Cre; Ndufs4^fl/fl^ animals. Note that the degeneration is stronger in the periphery of the optic nerve. B) Quantification of the number of axons in PPD-stained optic nerve sections. Multiple t-tests, \*\**P* < 0.01 and \*\*\**P* < 0.001).

**Suppl. Figure 3. Activated myeloid cells are present in the center and periphery of** α**-Cre; Ndufs4^fl/fl^ retinas.** Retinal flat-mounts from P45 α-Cre; Ndufs4^fl/fl^ mice immunostained for Iba1 (magenta) and Cd68 (teal) reveal a high density of activated myeloid cells across both central and peripheral regions of the GCL. Iba1_⁺_ cells co-express Cd68, a marker of activation and phagocytic activity, indicating widespread microglial activation in the degenerating retina.

**Suppl. Figure 4. Myeloid cell depletion improves survival of peripheral RGCs in** α**-Cre; Ndufs4^fl/fl^ retinas.** A) Retinal flat-mounts stained with RBPMS (teal) show a partial rescue of RGCs in the periphery of α*-*Cre; Ndufs4^fl/fl^ mice upon myeloid cell depletion with PLX5622 when compared to AIN76A control chow. B and C) Quantification of RBPMS+ RGCs per mm^2^ in animals fed with regular chow, AIN76A, or AIN76A supplemented with PLX5622. The central and peripheral retina were quantified separately (two-way ANOVA, n.s. not significant, \*\**P* < 0.01, \*\*\**P* < 0.001, and \*\*\*\**P* < 0.0001). ç

**Suppl. Figure 5. T-cells are present in** α**-Cre; Ndufs4^fl/fl^ retinas.** CD3 immunostaining (magenta) on flat-mounted retinas at P45 and P120 identified T-cells in both central and peripheral regions of α-Cre; Ndufs4^fl/fl^ (arrows). Note the unspecific labeling of erythrocytes within blood vessels in all conditions. All samples were counterstained with RBPMS to identify RGCs.

